## Supplementary text for "Galaxy-SynBioCAD: Automated Pipeline for Synthetic Biology Design and Engineering"

**Table of content**

SynBioCAD tools 2
Table S1. Galaxy-SynBioCAD tools 6
Tools design and integration process 8
Figure S1. Development and integration process. 9
Retrosynthesis workflow 10
Figure S2. RetroSynthesis and Pathway Enumeration workflow. 10
Pathway analysis workflow 11
Figure S3. Pathway Analysis and Ranking. 11
Figure S4. Top scored enumerated pathways. 13
Genetic design and engineering workflows 14
Golden Gate, Gibson and LCR assembly workflow 14
Figure S5. Genetic Design (Golden Gate, Gibson and LCR). 15
BASIC assembly workflow 15
Figure S6. BASIC assembly workflow. 16
Literature Pathways matching algorithm 17
Reaction matching score 17
Pathway matching score 18
Matching Threshold 18
Figure S7: Score distribution of all predicted pathways. 19
**References** 19

#### SynBioCAD tools

Tools described below are runnable as standalone tools in command-line terminal as well as Galaxy nodes through wrappers. Source code URLs and packaging details are given in Table S1.

**RetroRules**[1] is a searchable database of reaction rules. Because the target is likely not natively produced by the selected chassis, one cannot easily find a chemical reaction cascade that produces the target. A way to extend the capabilities description of the chassis is to generalize chemical reactions into reaction rules. RetroRules takes the space of known chemical reactions (roughly 40k reactions) and builds a set of generic reaction rules (about 350k). The RetroRules dataset provided presently is tagged as “rr02” and is freely downloadable from the RetroRules database at https://retrorules.org. The Galaxy RetroRules node provides an interface to directly download the RetroRules dataset, and to perform rule filtering based on (i) the rule diameter and (ii) the recommended rule usage. The node outputs a CSV file of reaction rules in SMARTS format.

**RetroPath2.0** is an open-source tool for building retrosynthesis networks by combining reaction rules and a retrosynthesis-based algorithm to link the desired target compound to a set of available precursors[2]. The RetroPath2.0 tool is freely available at myExperiment.org. In addition, a wrapper distributed as a conda package provides a convenient command line interface (CLI). The Galaxy RetroPath2.0 node takes as input two CSV files, one with a list of sink molecules using the standard InChI format, and a second file containing the reaction rules in SMARTS format. The target (source) molecule is provided using the InChI format through an input text box. The retrosynthesis network is outputted as a CSV file providing reactions in the reaction SMILES format and chemicals in both SMILES and InChI formats along with other information like the score for each reaction.

**RP2Paths** is an open-source tool dedicated to the enumeration of heterologous pathways that lie in a retrosynthesis network as produced by RetroPath2.0[2]. Such analysis is a required step in our workflow to ensure that only pathways fulfilling all the precursor needs are retained for further analysis. The node takes as input a retrosynthesis network in the CSV file produced by RetroPath2.0, and outputs the enumerated pathways (using IDs) as well as the structure of involved chemicals (as SMILES) in CSV files as well.

**rpCompletion** performs three essential steps, using as input the enumerated pathways from rp2paths and producing a collection of extended SBML files. First, because reaction rules are generic by design, each reaction rule can correspond to several template reactions: the task here is to enumerate the different possible transformations according to these templates. Then, because the RetroRules reaction rules consider only one substrate at a time, some compounds are by purpose omitted: the task here is to complete the predicted reactions by putting back these omitted compounds (mostly cofactors). Finally, the node converts each predicted pathway to distinct SBML files.

**rpThermo** calculates the Gibbs free energy of reactions and heterologous pathways by considering every chemical species involved in each reaction. This is done using the tool eQuilibrator[3] calculating the formation energy either using public database ID reference (when recognized with the tools internal database) or by decomposing the chemical structure and calculating its formation energy using the component contribution method. Thereafter, the species involved in a reaction are combined (with consideration for stoichiometry) and the thermodynamic feasibility of the pathway is estimated by taking the sum of the Gibbs free energy of each participating reaction. The node takes as input pathways in SBML format and returns annotated pathways (with thermodynamics information for each reaction, see Methods section for further details) also in SBML format.

**rpFBA** is used to calculate target production fluxes of the designed pathways. To perform Flux Balance Analysis (FBA) on a heterologous pathway, this tool first merges a heterologous pathway with a user-specified GEM model. The GEM models should be in the SBML format such as those available in the BiGG[4] and MetaNetX[5] databases. When merging the GEM model with heterologous pathways, compounds that cannot carry any flux are temporarily removed from the reaction for the FBA evaluation. Such cases can happen due to side substrates or products of predicted reactions that do not match any chassis compound. This enables FBA to consider whole-cell conditions for the theoretical production of the user's target molecule. The tool uses the COBRApy package to perform FBA[6]. The native COBRApy methods supported are FBA and parsimonious FBA (pFBA). The tool also contains an in-house developed method ("fraction of reaction" described in section Methods) to consider the potential burden that the production of a target molecule may have on the cell and the impact of the target itself. The node takes as input pathways like those produced by rp2Path and a strain model both in SBML format and returns annotated pathways (with calculated fluxes, see Methods section) in SBML format.

**rpScore** provides a global score for a given pathway. This score is computed by a machine learning (ML) model (see **Methods** section). The model takes as input features describing the pathway (thermodynamic feasibility, target flux with fixed biomass, length) and the reactions within the pathway (reaction SMARTS, Gibbs free energy, enzyme availability score) and prints out the probability for the pathway to be a valid pathway. The ML model has been trained on literature data (section Benchmarking with literature data) and by a validation trial (section Benchmarking by expert validation trial). Based on this global pathway score, a dedicated Galaxy wrapper (*rpRanker*) ranks a set of heterologous pathways to reveal what are the most likely pathways to produce the target molecule in an organism of choice.

**rpReport** generates HTML pages to visualize the main characteristics of the predicted pathways. Predicted pathways are summarized in a table, providing a quick overview of important characteristics, namely thermodynamics ($\Delta G'm$), fluxes (FBA for target production), the number of metabolic steps, reaction rule score (inherited from RetroRules rules), and the global score. Selecting one pathway shows information on its individual reactions using barplots, and lists of EC numbers with crosslinks. Selecting several pathways from the table render bar plots that compare these pathways. HTML reports are displayed using a responsive layout, it can be exported, and reports do not require an internet connection to be explored. The RP report node takes as input one or several pathways files in SBML format (.xml files), merged as a tar archive or in a folder. SBML files are parsed using python, and the HTML reports involve JavaScript code.

**rpViz** provides an interactive web interface for exploring predicted pathways and their associated annotations. The tool extracts information from the predicted pathways described as SBML files, and produces a HTML web page. The web page relies on JavaScript code and is a "dependency-free" output easy to set up locally for the user. Possible user interactions are pathway highlighting, cofactor handling, and the viewing of information at the levels of pathways, reactions, and involved compounds. The node takes as input pathways in SBML format.

**Selenzyme**[7] is an open-source tool that performs enzyme sequence selection from a reaction query. The tool can be queried using a reaction template such as the reaction rules in RetroRules. This feature makes this tool especially useful in combination with RetroPath2.0. Selenzyme performs a reaction similarity search in the reference reaction database Metanetx[5] and outputs the sequences annotated for the closest reactions. The tool provides several scores that can be combined in order to define an overall score. Scores are given for reaction similarity, conservation based on a multiple sequence alignment of the result, phylogenetic distance between source organism and host, and additional scores calculated from sequence properties. Selenzy takes input pathways in SBML format and returns annotated pathways (with UniProt ID for each reaction, see Method section) also in SBML format.

**SbmlToSbol** provides the mapping from the theoretical space to the practical space. This tool takes a pathway model (encoded in SBML) as input and returns a collection of placeholders for the subsequent design of the synthetic DNA that is required to encode the enzymes defined in the pathway model (encoded in SBOL). The converter first parses the SBML model and extracts a user-specified number of homologous enzymes for each metabolic reaction. Synthetic gene design templates, in the form of SBOL *ComponentDefinitions*, are generated for each enzyme, each consisting of an (enzyme) coding region (specified by a Uniprot sequence identifier), 5' and 3' flanking regions for downstream assembly, and - optionally - ribosome binding sites of user-specified translation initiation rates, allowing for the control of translational regulation. The SBOL document contains no sequence data but acts as a template to be passed onto the next node, PartsGenie.

**PartsGenie** is an established web application for the design of reusable synthetic DNA parts[8]. It supports the integrated design and optimization of ribosome binding sites, coding sequences, and other features, providing a multi-objective optimization algorithm that simultaneously optimizes translation initiation rate and codon usage along with elimination of repeating nucleotides and unwanted restriction sites. Furthermore, PartsGenie also implements guidelines from DNA manufacturers to optimize sequences for *synthesisability*, including the reduction of both local and global GC content. PartsGenie takes in the "template" SBOL document from the preceding SbmlToSbol converter step as input, and uses this a set of instructions to design and optimize synthetic DNA sequences for each gene in the template. The SBOL document is updated with these novel sequences. As PartsGenie is a REST service, a client has been developed to make requests.

**OptDoE** is based on the optimal design of experiments OptBioDes library[9] to combine selected genetic parts (from PartsGenie) and enzyme variants for the desired pathways from SBML files provided by Selenzyme. The *D*-optimal experimental design algorithm is based on a logistic regression analysis with an assumed linear model for the response evaluated based on its *D*-efficiency, which compares the design with an orthogonal design. The OptDoE node accepts as input the pathways in SBML format annotated with the enzyme variants and the collection of genetic parts consisting of plasmid copy numbers of the vector backbone, resistance cassette, promoters, and terminator in SBOL format and registered in the SynBioHub repository.

**DNA Weaver**[10] devises cloning strategies using either Golden Gate Assembly or Gibson Assembly to obtain plasmids for each combination of genetic parts selected by the OptDoE node. As both assembly methods have practical limitations, DNA weaver first considers Golden Gate assembly using the type-IIS enzymes BsmBI, BsaI, or BbsI (in this order) and defaults to Gibson Assembly, although this order of preference can be changed by the user. The resulting assembly strategies produce minimally "scarless" plasmids whose sequence is the direct concatenation of the sequences of the plasmid's parts, with the Golden Gate overlap sequences if this method is used. The node output is a spreadsheet featuring a list of all the primers required to extend the standard genetic parts with sequence homologies necessary for the assembly and a list of all PCRs and fragment assembly operations required to obtain the desired plasmids. The assembly strategy is optimized to maximize primer reuse between constructs and optimize assembly homologies, via the DNA Weaver framework[10]. A specific tool (DNAWeaver_SynbioCAD) has been written for SynBioCAD which takes SBOL designs produced by the SynbioCAD pipeline and device assembly plans based on this particular assembly options, and parameters and logics that are specific to SynbioCAD.

**LCR Genie**[11] is a web-based tool for supporting the design of bridging oligos, which are required for annealing together individual synthetic DNA parts (designed by PartsGenie) into multi-gene plasmid assemblies, designed by OptDoE. Promoters, RBSs and plasmid backbone are either chosen from a shortlist defined within the software (the default behavior) or provided by the user. Enzyme identifiers are randomly chosen and combined with the aforementioned parts to explore the combinatorics of possible constructs. LCR Genie provides a wrapper for this functionality, taking in an SBOL document containing numerous combinatorial plasmid assemblies, and designing bridging oligos necessary for assembly via the ligase cycling reaction method. The LCR Genie node performs analogous functionality to the DNA weaver node (supporting multi-part assembly but by a different experimental method) and as such, its output format matches that of DNA Weaver.

**rpBASICDesign** tool extracts enzyme IDs contained in SBML files — produced by tools such as Selenzyme — to generate genetic constructs compliant with the BASIC assembly approach[12]. The BASIC method relies on orthogonal linkers and type IIs restriction enzyme cleavage to provide a robust and accurate assembly of DNA parts into plasmid constructs. Different types of linkers (neutral, methylated Prefix-Suffix, RBS linkers) can be used for the assembly. Predefined sets of such linkers are commercially available as 96 well plates (*e.g.*, BioLegio plates[13]). rpBASICDesign uses as input an SBML file annotated with enzyme IDs for each reaction, and optionally one or several files listing by their IDs the linkers, the promoters and the backbone used. If not provided, then a default list of parts is used. It produces 3 CSVs and a set of SBOL files. The main one lists the constructs to be built, where each construct is described by a row and consists of a sequence of BASIC linker and DNA part IDs. The 2 other CSV files provide the plate coordinates of the BASIC linkers and the DNA-parts that the user will need to provide. Additionally, one SBOL file is produced for each construct generated. For a given set of enzyme coding genes as standardized DNA parts, several combinations of promoters and RBSs are generated, and permutations of the gene order can be optionally performed in an operon format.

**DNA-Bot** software takes as input the CSV files describing the construct (such as those produced by rpBASICDesign) and generates instructions for the automated build of the genetic constructs using OpenTrons liquid handling robots. Optional parameters can be set by the user to define the plastic labwares to be used, and set protocol parameters such as washing or incubation times for purification step. DNA-Bot outputs python scripts that implement the 4 assembly steps, namely Clip reactions, Purification, Assembly, and Transformation. In short, the Clip reactions step prepares the mixes for the ligation of the individual DNA parts with the linkers; the Purification step purifies the linker-ligated DNA parts using magnetic beads and the Opentrons magnetic module; the Assembly step mixes the DNA purified parts to build the final constructs; while the Transformation step transforms the chassis micro-organism with the plasmid and inoculates onto agar. Additional metadata meaningful to keep track of parameters are also outputted by the tool.

The Galaxy-SynBioCAD portal does not currently support the visualization of SBOL files such as those produced by PartsGenie and OptDoE, however, these files can be downloaded and visualized using online tools such as VisBOL[14]. Alternatively, they can be uploaded to an SBOL repository and viewed with SBOL visual representation and the newly incorporated sequence viewer.

The Galaxy-SynBioCAD portal also supports other nodes not listed above that perform simple operations like uploading a file, extracting taxonomy ID, or native metabolites from Genome-scale metabolic models (GEMs) file (SBML).

**Table S1. Galaxy-SynBioCAD tools**

| NAME | SOURCE CODE | CONDA package channel | GALAXY WRAPPER | GALAXY NODE |
|---|---|---|---|---|
| RRParser | https://github/brsynth/RRParser | rrparser conda-forge | https://github.com/brsynth/synbiocad-galaxy-wrappers/tree/master/RRulesParser | https://toolshed.g2.bx.psu.edu/view/tduigou/rrparser/ea590c609fec |
| rpExtractSink | https://github.com/brsynth/rptools/tree/master/rptools/rpcompletion | rptools conda-forge | https://github.com/brsynth/synbiocad-galaxy-wrappers/tree/master/rpExtractSink | https://toolshed.g2.bx.psu.edu/view/tduigou/rpextractsink/47bb93e7832b |
| RetroPath2.0 | https://github.com/brsynth/RetroPath2-wrapper | retropath2_wrapper | https://github.com/brsynth/synbiocad-galaxy- | https://toolshed.g2.bx.psu.edu/view/tduigou/retropath2/9c8ac |

| NAME | SOURCE CODE | CONDA package channel | GALAXY WRAPPER | GALAXY NODE |
|---|---|---|---|---|
| | | conda-forge | wrappers/tree/master/RetroPath2-wrapper | 9980bd6 |
| RP2Paths | https://github.com/brsynth/rp2paths | rp2paths conda-forge | https://github.com/galaxy-synbiocad/galaxy-nodes/tree/master/rp2paths | https://toolshed.g2.bx.psu.edu/view/tduigou/rp2paths/2782bee7c5a6 |
| rpCompletion | https://github.com/brsynth/rptools/tree/master/rptools/rpcompletion | rptools conda-forge | https://github.com/brsynth/synbiocad-galaxy-wrappers/tree/master/rpCompletion | https://toolshed.g2.bx.psu.edu/view/tduigou/rpcompletion/b8242cf18cc0 |
| rpThermo | https://github.com/brsynth/rptools/tree/master/rptools/rpthermo | rptools conda-forge | https://github.com/brsynth/synbiocad-galaxy-wrappers/tree/master/rpThermo | https://toolshed.g2.bx.psu.edu/view/tduigou/rpthermo/21a900eee812 |
| rpFBA | https://github.com/brsynth/rptools/tree/master/rptools/rpfba | rptools conda-forge | https://github.com/brsynth/synbiocad-galaxy-wrappers/tree/master/rpFBA | https://toolshed.g2.bx.psu.edu/view/tduigou/rpfba/c554f15279fe |
| rpScore | https://github.com/brsynth/rptools/tree/master/rptools/rpscore | rptools conda-forge | https://github.com/brsynth/synbiocad-galaxy-wrappers/tree/master/rpscore | https://toolshed.g2.bx.psu.edu/view/tduigou/rpscore/da8ae7fa5ed3 |
| rpRank | https://github.com/brsynth/rptools/tree/master/rptools/rprank | rptools conda-forge | https://github.com/brsynth/synbiocad-galaxy-wrappers/tree/master/rpRanker | https://toolshed.g2.bx.psu.edu/view/tduigou/rpranker/e95370d2e5f9 |
| rpReport | https://github.com/brsynth/rptools/tree/master/rptools/rpreport | rptools conda-forge | https://github.com/brsynth/synbiocad-galaxy-wrappers/tree/master/rpreport | https://toolshed.g2.bx.psu.edu/view/tduigou/rpreport/d09a51507aaf |
| rpViz | https://github.com/brsynth/rptools/tree/master/rptools/rpviz | rptools conda-forge | https://github.com/brsynth/synbiocad-galaxy-wrappers/tree/master/rpviz | https://toolshed.g2.bx.psu.edu/view/tduigou/rpviz/ea2ca40a24c5 |
| Selenzy | https://github.com/pablocarb/selenzy | selenzy_wrapper bioconda | https://github.com/brsynth/synbiocad-galaxy-wrappers/tree/master/selenzy-wrapper | https://toolshed.g2.bx.psu.edu/view/tduigou/selenzy/34a9d136a5bf |
| SbmlToSbol | https://github.com/neilswainston/SbmlToSbol | sbml2sbol conda-forge | https://github.com/brsynth/synbiocad-galaxy-wrappers/tree/master/sbml2sbol | https://toolshed.g2.bx.psu.edu/view/tduigou/sbml2sbol/83108f3c65aa |
| PartsGenie | https://github.com/neilswainston/PartsGenie | | | |
| PartsGenie Client | https://github.com/neilswainston/PartsGenieClient | | https://github.com/brsynth/synbiocad-galaxy-wrappers/tree/master/PartsGenie | https://toolshed.g2.bx.psu.edu/view/tduigou/partsgenie/295a21fc55d0 |

| NAME | SOURCE CODE | **CONDA** package channel | GALAXY WRAPPER | GALAXY NODE |
|---|---|---|---|---|
| OptDoE | https://github.com/pablocarb/doebase | doebase conda-forge | https://anaconda.org/conda-forge/doebase | https://toolshed.g2.bx.psu.edu/view/tduigou/optdoe/c3f32929a4b7 |
| DNAWeaver | https://github.com/Edinburgh-Genome-Foundry/DnaWeaver | dnaweaver bioconda | | |
| DNAWeaver SynBioCAD | https://github.com/brsynth/DNAWeaver_SynBioCAD | dnaweaver_synbiocad bioconda | https://github.com/brsynth/synbiocad-galaxy-wrappers/tree/master/DNAWeaver | https://toolshed.g2.bx.psu.edu/view/tduigou/dnaweaver/c519517e3ade |
| LCRGenie | https://github.com/neilswainston/LCRGenie | lcr_genie conda-forge | https://anaconda.org/conda-forge/lcr_genie | https://toolshed.g2.bx.psu.edu/view/tduigou/lcrgenie/afbbecdcc0e3 |
| rpBASICDesign | https://github.com/brsynth/rpbasicdesign | rpbasicdesign conda-forge | https://anaconda.org/conda-forge/rpbasicdesign | https://toolshed.g2.bx.psu.edu/view/tduigou/rpbasicdesign/de9f53630349 |
| DNA-Bot | https://github.com/BASIC-DNA-ASSEMBLY/DNA-BOT | dnabot conda-forge | https://anaconda.org/conda-forge/dnabot | https://toolshed.g2.bx.psu.edu/view/tduigou/dnabot/6d55c77a17ab |

### Tools design and integration process

The above tools have been put together within a unified and user-friendly interface (Galaxy). As shown in **Figure S1** below, they have been developed as standalone command-line tools without graphical interface and, to be fully compliant with Galaxy packages management, published as Conda packages on Anaconda.org[15]. These packages run mostly on all major Linux distributions, MacOS and Windows and are downloaded from remote channels (conda-forge or bioconda). During the development process, source code changes trigger GitHub actions, a continuous integration platform which helps to automatically run the test processes and displays the build status. We also designed and developed wrappers to make tools available through Galaxy which automatically download the previous packaged tools from anaconda in order to install all the needed dependencies.

Each wrapper is deposited in its own GitHub repository, isolated from the core source code of the corresponding tool. The wrappers are tested using Planemo Utilities[16], which helps to check the XML validity, to run the tests already written in the test section and finally to publish all the tools to a test ToolShed[17] and the main ToolShed[18].

Concerning the development process, we have followed the FAIR (Findability, Accessibility, Interoperability, Reproducibility) principles. Thus, (i) source codes are freely accessible on public repositories, (ii) each tool is published as a package installable on main OS platforms (reusable in space) with versioning capabilities (reusable in time), and (iii) standard formats are used as tools input and output to ensure connections with the outside of the current ecosystem. To go further in FAIR principles, each tool is also available within the Galaxy scientific workflow manager whose key principles are precisely accessibility, transparency and reproducibility of workflows. The Galaxy system provides graphical user interfaces to combine different technologies along with efficient methods for using, sharing and publishing them and thus increasing the efficiency of the scientists using them.

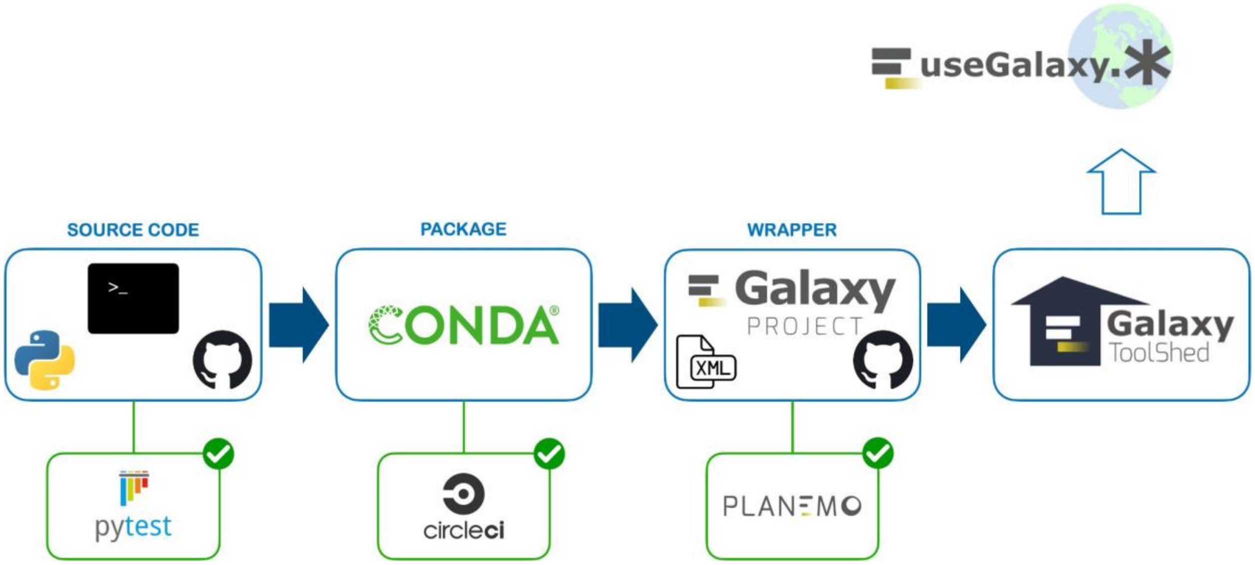


**Figure S1**. **Development and integration process**.

*Tools are written in Python and runnable in a command-line terminal, source code is versioned on GitHub. Tests are processed with pytest before releasing a new version. Then, tools are packaged under Conda format, tested and published on anaconda.org. Galaxy wrappers are written in XML to make tools available (after tests validation with CircleCI) within Galaxy. Finally, tools are published in the Galaxy ToolShed using Planemo utilities to be available on Galaxy community portals.*

### Retrosynthesis workflow

The first workflow, illustrated in **Figure S2**, is the one that processes retrosynthesis and pathway enumeration. The workflow takes as input: (i) the International Chemical Identifier (InChI) of the compound of interest to produce, (ii) the SMBL genome scale model of the chosen chassis organism, and (iii) the reaction rules (generated by RRules Parser node that calls RetroRules).

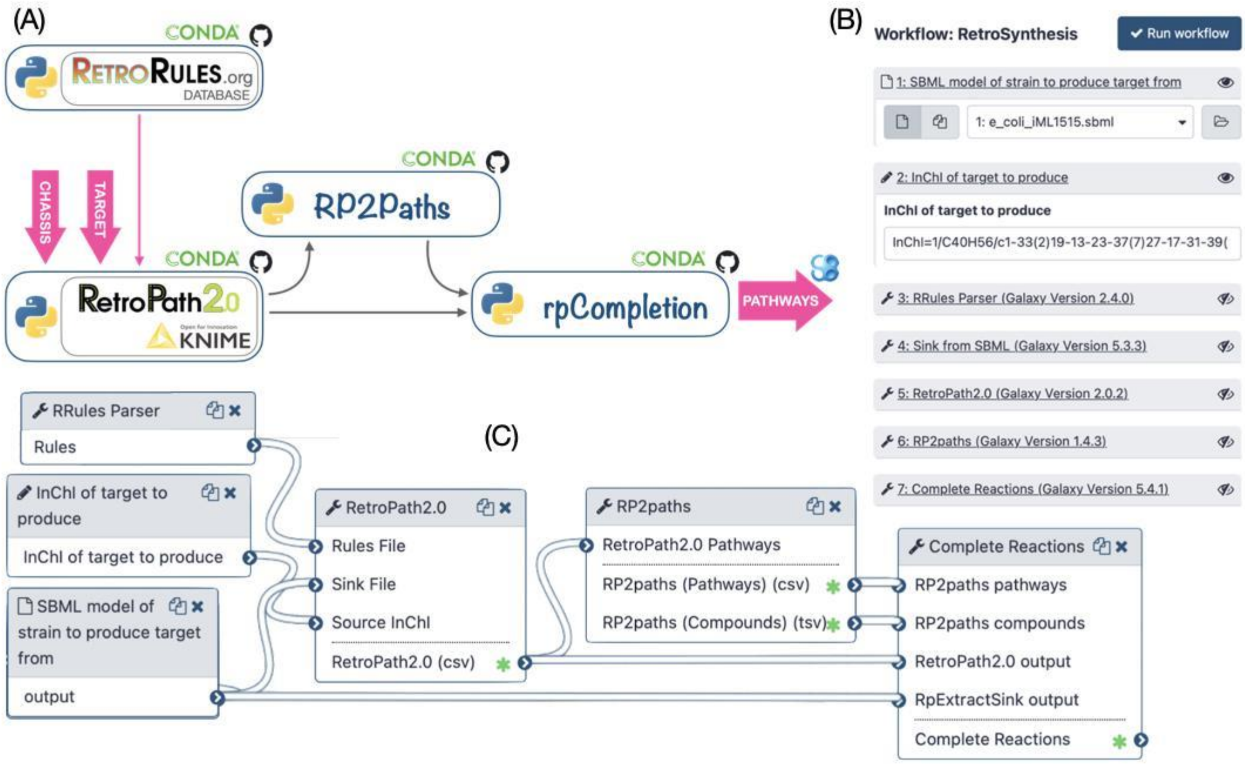


**Figure S2. RetroSynthesis and Pathway Enumeration workflow.**
***(A)*** *The workflow of tools for retrosynthesis and pathway enumeration. Tools can be chained manually by running each tool one after the other in a command-line terminal. Outputs of each tool can be directly given as inputs of the others without any other processing.* ***(B)*** *The workflow menu at runtime in the Galaxy interface. The user specifies the genome scale SBML model of the host organism and the InChI structure of the target molecule. The user can also change the default settings for each tool by clicking on its name. The RetroRules entry has been set as default for convenience. The workflow generates a collection of heterologous pathways for target production in separate SBML files.* ***(C)*** *The workflow as displayed in the Galaxy workflow Editor.*

The workflow generates theoretical possible pathways for the production of a target molecule in a chosen organism. Three key steps are performed in this workflow, these are detailed in the Methods section (*cf.* Reaction rules, Retrosynthesis from target to sink, Pathway annotation) and summarized next. First, using RetroPath2.0, the workflow generates a network of feasible metabolic routes to produce a target molecule in a selected chassis organism. That metabolic network is then decomposed into individual pathways using RP2paths. Lastly, rpCompletion takes those individual metabolic pathways to filter them (duplicated pathways are removed), then splits them into sub-pathways by adding the appropriate cofactors, and finally converted them to SBML files. Additional details are provided in the Methods section (*cf*. Pathway completion combinatorics)

#### Pathway analysis workflow

This workflow ranks a set of pathways based on multiple metrics (flux balance analysis, thermodynamics, pathway length, and reaction SMARTS, *cf*. **Figure S3**). The workflow takes as inputs: (i) the list of pathways to rank, and (ii) the structure of metabolites present in the chosen chassis organism (e.g., *E. coli* model iML1515).

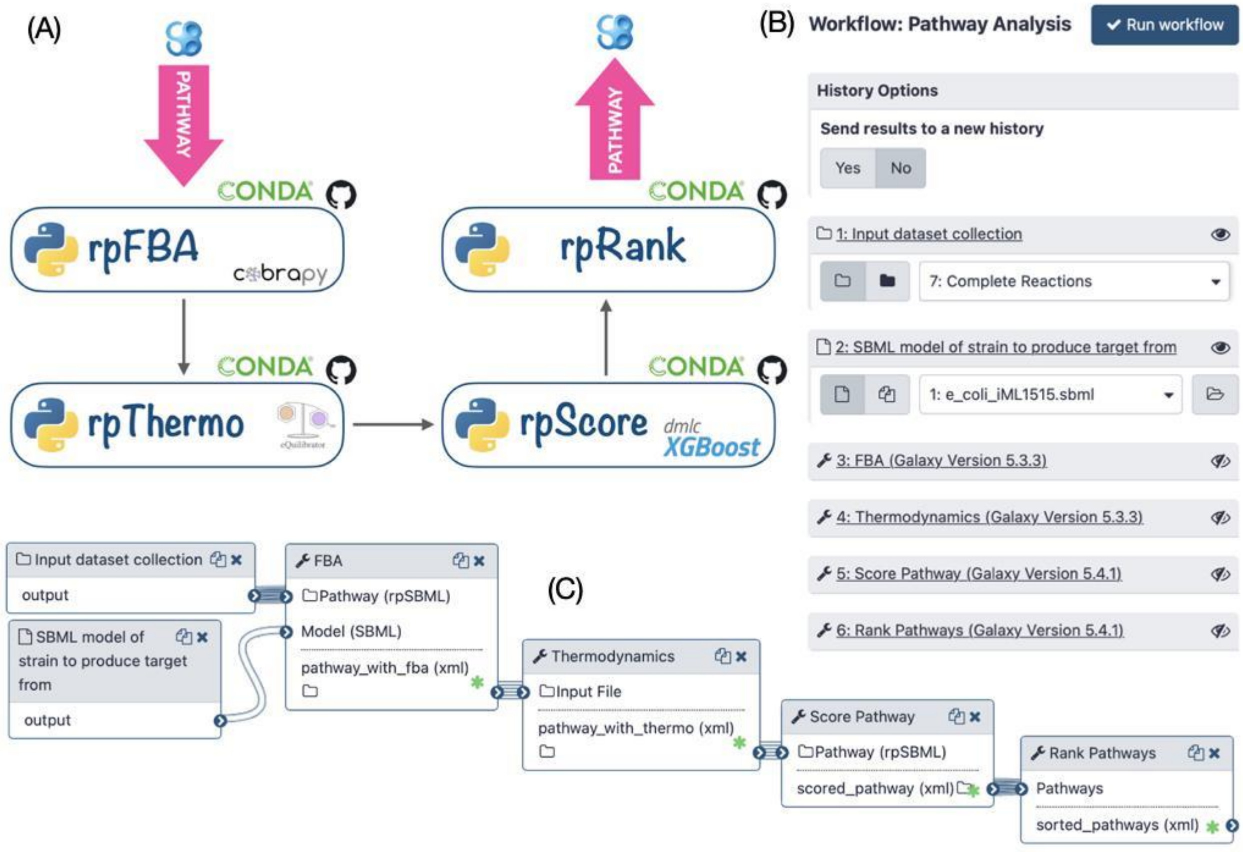


**Figure S3. Pathway Analysis and Ranking.**

***(A)** Workflow of tools for pathway analysis and ranking. **(B)** The workflow menu upon executing it within the Galaxy interface. The user specifies the GEM SBML model of the host organism and the set of pathways to rank (here we can choose the output of Retrosynthesis workflow). The user can also modify default parameters of each tool by clicking on its name. The workflow generates a collection of heterologous pathways which are scored and ranked. **(C)** The workflow as displayed in the Galaxy workflow manager.*

Given a set of pathways generated by the *Retrosynthesis* workflow, *Pathway analysis* informs the user as to the theoretically best performing taking various criteria calculated by the rpScore tool. The criteria used for scoring are listed below.

1. In the *Retrosynthesis* workflow, molecules contained within a full SBML model are used to compute heterologous pathways. As a result, the calculated heterologous pathways can easily be merged into the full organism model, enabling the whole-cell context to calculate the production flux of a given target. The method forces a fraction of its maximal flux through the biomass reaction while optimizing for the target molecule. This is achieved through the FBA node. The FBA node is further described in **Methods** (*cf*. **Flux Balance Analysis with Fraction of Reaction**).
2. Thermodynamics values (based on Gibbs free energies) are computed for each pathway by using a linear equation system solver (*cf.* T**hermodynamics** in **Methods**) to optimize the yield of the reaction producing the target and to remove intermediate compounds to not clutter up the cell.
3. Enzyme availability for the chemical transformation is also taken into consideration, where high values favor less promiscuous reaction rules and express better confidence. The method used to compute enzyme availability score is described the **Methods** section (*cf*. **Retrosynthesis from target to chassis**).
4. Finally, the length of the pathway is taken into consideration, here shorter pathways are favored over longer pathways.

Lastly, the above metrics are given to a machine learning model (*cf.* **Machine Learning Global Scoring** in **Methods**) in prediction mode to provide a single global score per pathway. The results may be graphically inspected by the user using a Galaxy embedded visualizer (rpViz node). The visualizer displays the heterologous metabolic routes, where complete descriptions of the chemical species, reaction and pathways are displayed. **Figure S4** presents displays from rpViz and rpReport for production of Lycopene in *E. coli* (model iML1515).

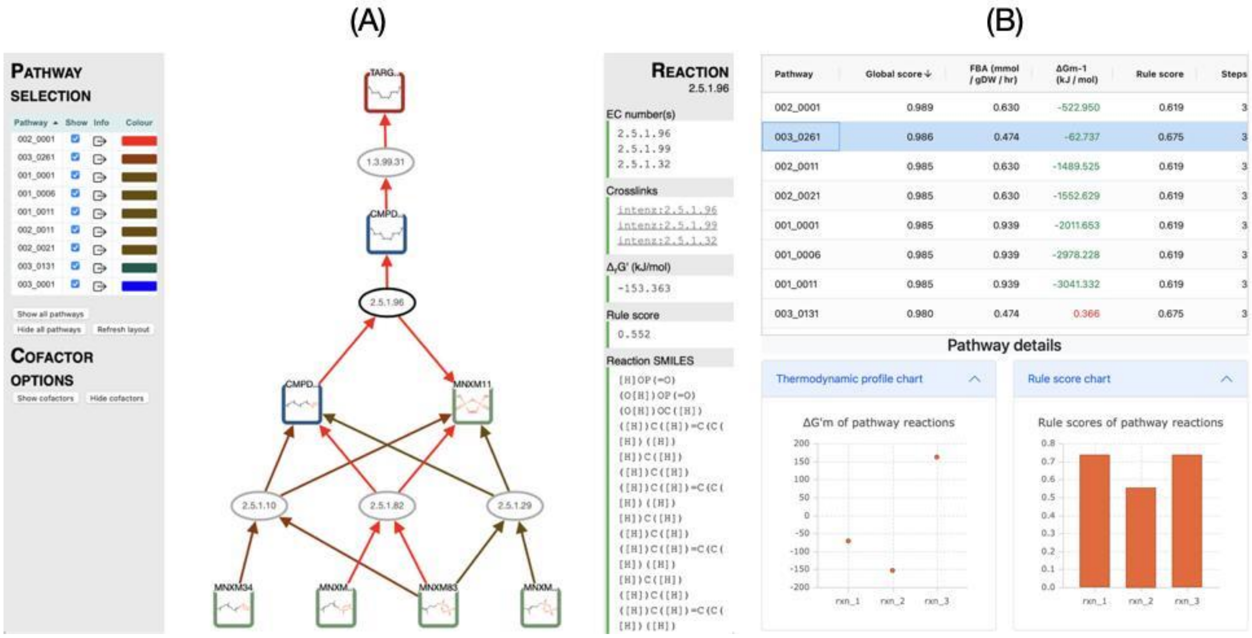


#### Figure S4. Top scored enumerated pathways.

*The example plots the top ranking pathways for the production of lycopene in E. coli after running the Pathway Analysis workflow.* ***(A)*** *In rpViz, the squares depict the molecules and the ovals the reactions. The green squares are the compounds that exist in the genome scale model (here iML1515), the blue squares are intermediate compounds, and the red square is the target. Each compound and reaction can be selected and the right-hand side displays details of the selection (here for the reaction 2.5.1.96). The panel also offers a link to look up the reaction on the Selenzyme web service to manually search enzymes that may perform such chemical transformations. On the left-hand side is the ranked list of pathways predicted, color-coded so that the best theoretical performing ones have warmer colors. The user may inspect the pathway as a whole by selecting the boxed arrow. This action displays on the right-hand panel information on the pathway including the number of steps, its thermodynamic feasibility, its flux, and its global score. The user can also display the cofactors for all the reactions by selecting the "Show cofactors" button on the left side panel.* ***(B)*** *This part is produced by the tool rpReport, the top-panel displays the pathways with for each global score, FBA, thermodynamics, rule score and number of steps. By clicking on one category, pathways will be ranked by this category (here by global score). After selecting one pathway (here 003_0261), the user can visualize pathway details on the bottom-panel including thermodynamic values (on the left) and rule score (on the right). The user can also select all pathways on the top-panel to have comparative charts.*

### Genetic design and engineering workflows

The previous workflow (*Pathway Analysis)* provides a list of ranked metabolic pathways producing a molecule of interest within a selected chassis organism. The next step is to engineer all or some of these pathways. Pathway selection can be performed in a fully automatic way, by retaining for instance the top ranked pathway. Reviewing the pathways using rpViz is also a good option to let experts browse prediction and select the best implementations. Once pathways have been selected, we provide two genetic design workflows for different assembly protocols. The first workflow provides assembly plans by using three different techniques: Golden Gate[19], Gibson[20], and Ligation Chain Reaction (LCR)[21]. The workflow offers the possibility of using the OptDoE node for combinatorial experimental design. We also provide a second workflow using Biopart Assembly Standard for Idempotent Cloning (BASIC) technique[12]. This workflow takes as input a pathway (in SBML format) and generates a script to operate an Opentrons liquid handler robot which performs assembly and chassis transformation.

#### *Golden Gate, Gibson and LCR assembly workflow*

This workflow, illustrated in **Figure S5**, encodes the top-ranking predicted pathways from the *Pathway Analysis* workflow into plasmids intended to be expressed in the specified organism. First, the Selenzyme node is executed to return a user-defined number of UniProt ID's associated with each reaction. Then a maximum number of pathways, defined by the user, are converted to an SBOL file. The next tool, PartsGenie, then retrieves the DNA sequences of the predicted enzymes based on their Uniprot ID, performs a codon optimization and creates a first level of library based on those, adding before the CDS some specific strength calculated RBS; these sequences can be output for direct gene synthesis. These parts are then used by OptDoE to design a defined size library of plasmids, expressing at various levels the genes coding for the multiple enzymes present in the predicted pathways. The other genetic parts required by this software (origin of replications, promoters, terminators and markers) are either provided by a default list or a specific list of parts provided by the user which needs to refer to parts stored in SynBioHub. The Galaxy tool "OptDoE Parts Reference Generator" has been written for that purpose. This final genetic design library is generated in an SBOL format and can then be used as an input to other softwares or visualized using tools implementing the SBOL visual standard. The *Golden Gate & Gibson assembly* workflow ends with two different tools tackling the library construction problem: LCR Genie that proposes an assembly strategy using the Ligase Chain Reaction method and DNA weaver, which calculates the optimal synthesis plan and the assembly protocol following either a Golden Gate or a Gibson Assembly method. The output of LCR Genie or DNA weaver is an excel file containing the full sequences of the plasmid library and of the intermediate parts required to construct them.

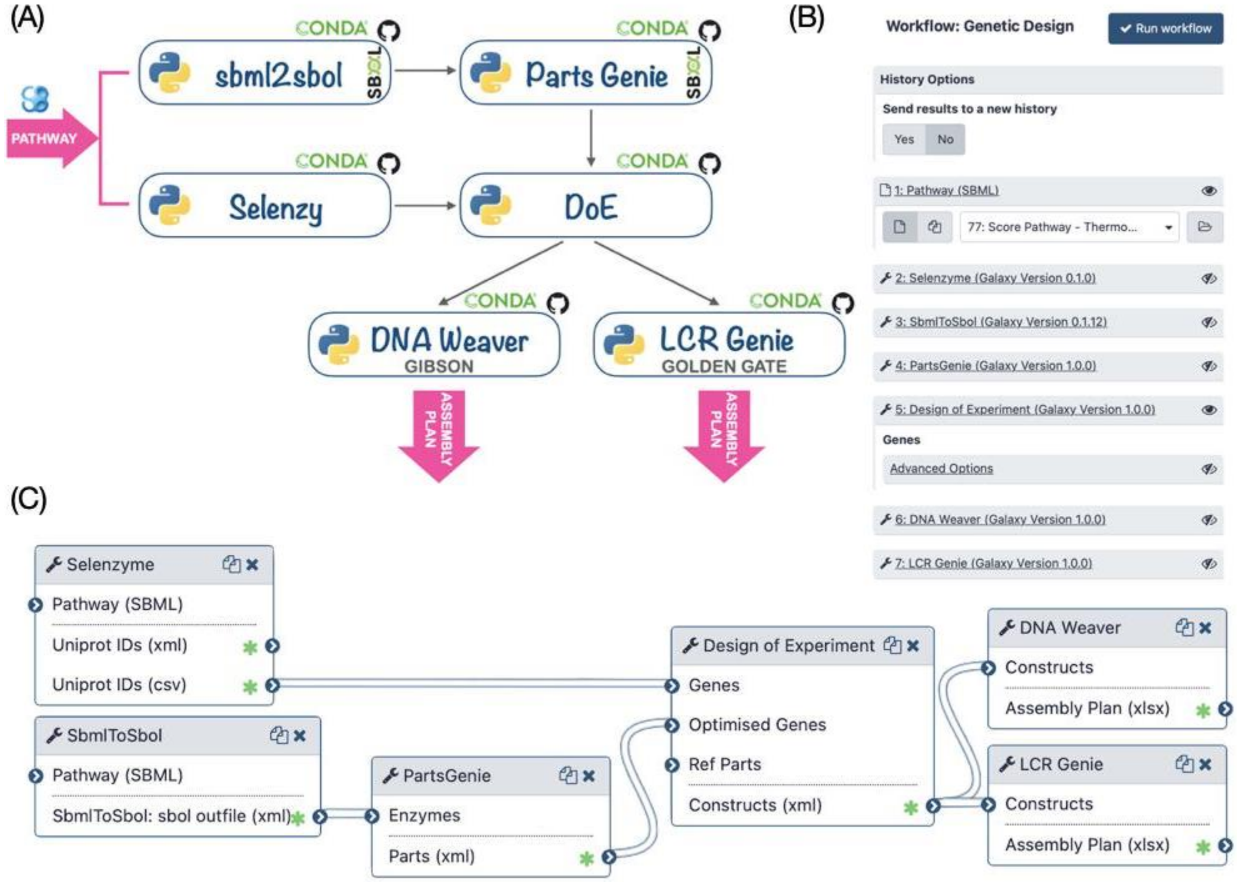


**Figure S5. Genetic Design (Golden Gate, Gibson and LCR).**

***(A)*** *Tools can be chained manually by running each tool one after the other in a command-line terminal. Outputs of each tool can be directly given as inputs of the others without any other processing.* ***(B)*** *The workflow menu upon executing it through the Galaxy interface. The user specifies the pathway (SBML) that he wishes to build. The workflow generates assembly plans by using LCR (LCR Genie) or Golden Gate or Gibson (DNA Weaver).* ***(C)*** *The workflow as displayed in the Galaxy workflow manager.*

#### *BASIC assembly workflow*

The Galaxy workflow is depicted in **Figure S6.A** and further illustrated in the Methods section (*cf*. **Basic Design and DNA-BOT workflow execution**). At first, a pathway generated by either the *RetroSynthesis* or the *Pathway Analysis* workflows is provided as an SBML file to the Selenzyme tool. Selenzyme searches for enzymes corresponding to each reaction of the pathway, and outputs an updated SBML file annotated with the enzyme UniProt IDs. To restrict the enzyme search to only a subpart of the tree of life (*e.g*. only enterobacteria) a list of taxonomic IDs can be provided. Second, the BasicDesign tool converts the SBML file into CSV files describing the DNA-parts to be included into each construct (in an operon format). Depending on the numbers of enzymes per reaction, of RBSs and promoters available, and whether or not to perform CDS permutation within the operon, the number of constructs may vary. In the last step, the DNA-Bot tool reads the list of constructs and the DNA-parts position on the source plates and generates a set of python scripts to build the plasmids using an Opentrons liquid handling robot. After downloading these scripts onto a computer connected to an Opentrons, the user can perform the automated construction of the plasmids at the bench.

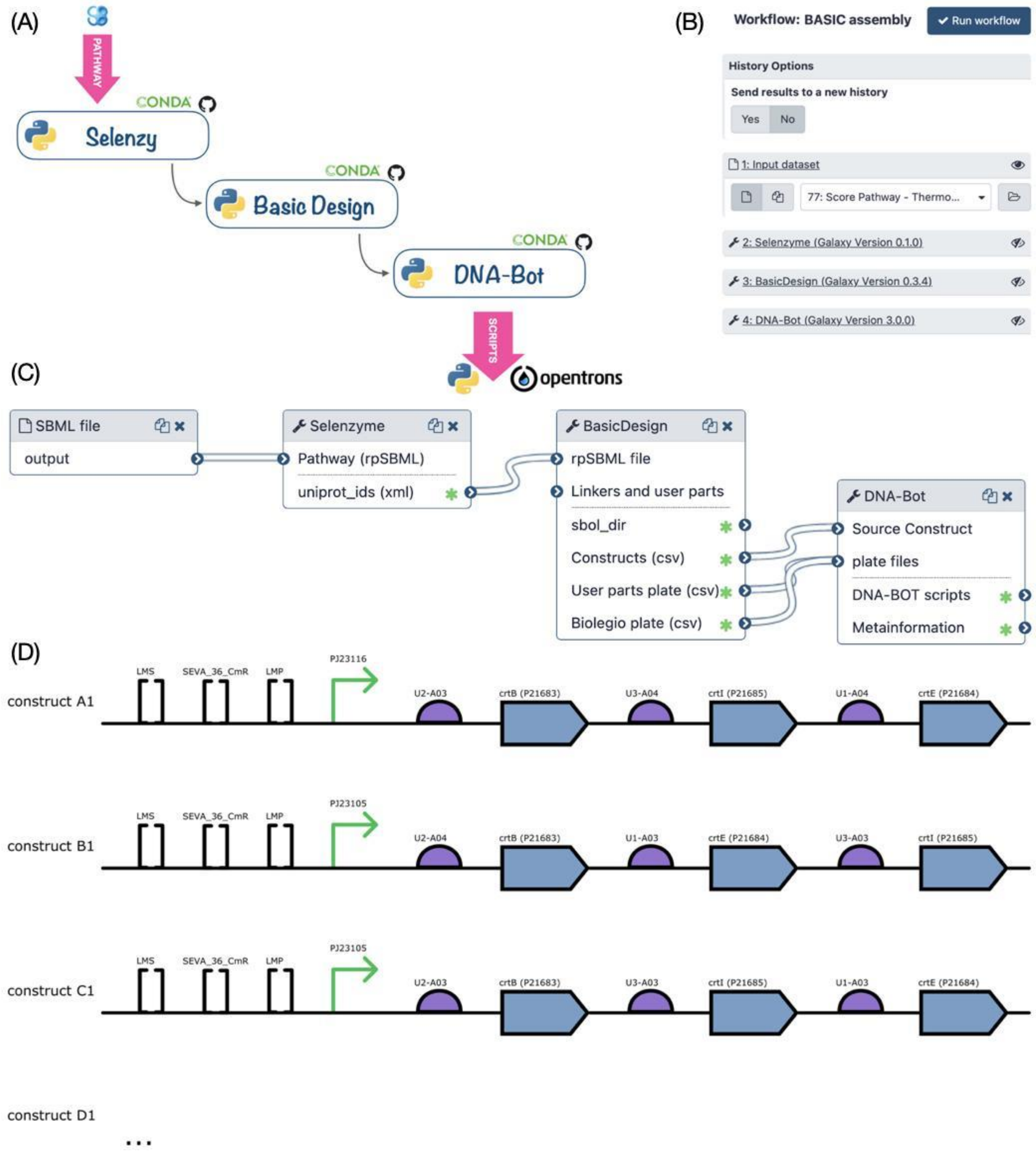


**Figure S6. BASIC assembly workflow.**

***(A)*** *The three genetic design tools can be chained manually by running each tool one after the other in a command-line terminal. Outputs of each tool can be directly given as inputs of the others without any other processing.* ***(B)*** *The workflow menu upon executing it through the Galaxy interface. The user specifies the pathway (SBML) that he* *wishes to build.* ***(C)*** *The workflow as displayed in the Galaxy workflow manager.* ***(D)*** *Architecture of the three first constructs generated by BasicDesign for the lycopene pathway in SBOL format. The view representation is generated using the VisBOL web service. Squared brackets represent "miscellaneous" parts corresponding to methylated prefix and suffix linkers (LMS and LMP) and the plasmid backbone (BASIC_S...). Other parts (promoter, RBS, CDS) are shown using the usual SBOL symbols, the RBS sequences are coded on standardized UTR-RBS linkers and so form the linkers between the promoter and CDS parts.*

### Literature Pathways matching algorithm

An algorithm was designed to quantify the degree of similarity between a *true* pathway (from the literature) and a list of *predicted* pathways generated by the Galaxy-SynBioCAD. To test the algorithm, we compiled 79 pathways from literature (cf. Supplementary file 'Literature_Pathways') and used the *Retrosynthesis* and *Pathway Analysis* workflows to generate *predicted* pathways for the same targets and chassis organisms. Since extracting information from journal articles can be difficult and reports are commonly incomplete, the algorithm reports to what degree of confidence a *predicted* pathway matches a *true* one. Let us note that the *predicted* pathway contains all the necessary information while the *true* pathway may only contain partial information. Pathways are first compared at the reaction level (all reactions from the one that generated the target to the one that is linked to the chassis organism) then at the pathway level.

#### *Reaction matching score*

To compare two reactions $r_{true}$ and $r_{pred}$, all the reactants of the *true* reaction are compared with all the reactants of the *predicted* reaction, the same comparison is performed for the products. When several reactants (products) are present in $r_{true}$, one searches in $r_{pred}$, the most similar reactant (product). Similarity between two given *true* and *pred* reactants (products) is carried out using the Morgan fingerprints of the reactants (products) computed using the RDKit library and a Jaccard coefficient (also named Tanimoto coefficient) is calculated from the fingerprints. The chemical reaction score $R_{CH}$ is the averaged Jaccard coefficient computed for all the reactants and the products of $r_{true}$.

The second criteria ($R_{EC}$ score) to match reactions is based on the EC numbers of the *true* and *predicted* reactions up to the fourth level. If the two reactions have the same first digit EC number then 1/2 is added to $R_{EC}$, if the reactions then match as the second level 1/4 is added, 3/16 is added if the reactions match at the third level, and 1/16 at the fourth level. As $R_{CH}$, $R_{EC}$ ranges from 0 to 1.

Lastly, the two scores are combined with a weighted mean:

$$r = (0.8, 0.2)\ (R_{CH}, R_{EC})^{T} \qquad (1)$$

Similarly to the species match, a measured reaction can match multiple predicted ones and thus the matches are computed in a matrix and the algorithm selects the best one.

#### *Pathway matching score*

Because predicted pathways can be of different lengths to the measured one, we define a pathway length penalty score:

$$h = 1.0 - \frac{|length(true) - length(pred)|}{max\ (\ length(true), length(pred)\ )} \quad (2)$$

Obviously h = 1.0 when the two pathways (*true, pred*) have the same number of steps. The penalty is applied to the sum of the reaction match score giving the final pathway match score:

$$f = h\,\overline{r_m} \quad (3)$$

where $\overline{r_m}$ is the mean score over all the *m* reactions in the pathway.

#### *Matching Threshold*

To determine a matching score threshold, above which a *predicted* pathway can be considered identical to a *true* pathway, we collected for each literature pathway the *predicted* pathway having the best matching score using eq. (3). Pathways can differ because they use different enzymatic reactions with different substrates and products. While the *predicted* pathways generated by SynBioCAD contain all the substrates and products of the reactions, cofactors (cosubstrates and coproducts) are generally not reported in the literature and matching score can be lowered because of these missing cofactors. To verify that the main substrate and main product of a given *true* pathway are accounted for in a *predicted* pathway, we compute a similarity (using Jaccard coefficient) between the two pathways removing cofactors. Precisely, for each reaction of the *true* pathway a chemical reaction score ($R_{CH}$ as defined in eq. 1) is computed without taking cofactor into account, in other words one searches in the predicted pathways the substrate (product) most similar to the one found in the *true* reaction. The similarity between the *true* and *predicted* pathway is the average $R_{CH}$ value for each *true* reaction. A Jaccard coefficient of 1 indicates that the main substrates and products for all reactions of a literature pathway are retrieved in the predicted pathway.

One observes from **Figure 2.D** (main text) that when the matching score is above 0.5 the predicted reactions are nearly identical to the literature reactions (with a pathway similarity of 1). Consequently, any pathway generated by Galaxy-SynBioCAD is labeled ‘literature pathway’ if its score is above 0.5. **Figure S7** below gives the distribution of matching scores obtained for all generated pathways, about 20% are passing the threshold constraint.

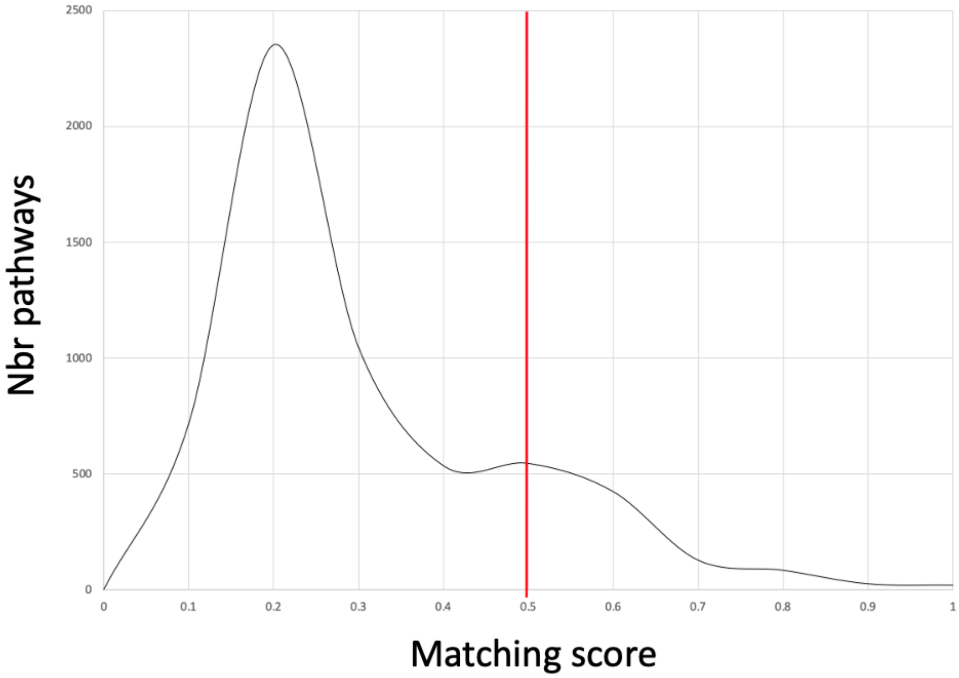


**Figure S7: Score distribution of all predicted pathways.**
The total number of pathways generated is 5874, 1222 (~20%) of which have a score above 0.5.

13. BASIC DNA Assembly | Biolegio. https://www.biolegio.com/products-services/basic/

14. VisBOL Design Visualizer. https://visbol.org

15. Anaconda.org. https://anaconda.org/

16. Planemo — Planemo's documentation. https://planemo.readthedocs.io/en/latest/

17. Galaxy Test Tool Shed. https://testtoolshed.g2.bx.psu.edu/

18. Galaxy Main Tool Shed. https://toolshed.g2.bx.psu.edu/

19. Engler, C., Gruetzner, R., Kandzia, R. & Marillonnet, S. Golden gate shuffling: a one-pot DNA shuffling method based on type IIs restriction enzymes. *PloS one* **4**, e5553 (2009).

20. Gibson, D. G. *et al.* Enzymatic assembly of DNA molecules up to several hundred kilobases. *Nature methods* **6**, 343–5 (2009).

21. Wiedmann, M. *et al.* Ligase chain reaction (LCR)--overview and applications. *Genome Res.* **3**, S51–S64 (1994).
